# Risk of Infection Due to Airborne Virus in Classroom Environments Lacking Mechanical Ventilation

**DOI:** 10.1101/2022.12.15.520644

**Authors:** Alexandra Goldblatt, Michael J. Loccisano, Mazharul I. Mahe, John J. Dennehy, Fabrizio Spagnolo

**Author notes:** Authors contributed equally (Alexandra Goldblatt, Michael J. Loccisano, Mazharul I. Mahe). Funding: This work was funded by NSF RAPID Award No. 2032634 (JJD).

## Abstract

The COVID-19 pandemic highlighted the role of indoor environments on disease transmission. Enclosed spaces where pathogen-laden aerosols accumulate was strongly linked to increased transmission events. Here we employ a surrogate non-pathogenic virus, the bacteriophage phi6, to interrogate aerosol transmission in classroom environments that do not have any natural or mechanical ventilation in order to determine how effectively aerosols facilitate new infections. We find that virus-laden aerosols establish new infections over all distances tested within minutes and that the time of exposure did not change transmission rate. We further find that humidity, but not temperature nor a UV-based disinfection device, significantly impacted transmission rates. Our data suggest that, even without mechanical ventilation, relative humidity remains a highly effective mitigation strategy while UV air treatment did not.

**Practical Implications:** Transmission of pathogens through airborne particles is a major source of disease transmission. People now spend much of their time indoors, thus understanding indoor airborne transmission is vital to managing outbreaks. Most classrooms in the U.S. do not have any mechanical ventilation systems and so here, we test airborne transmission of a virus in such classrooms. Infection transmission rates are not greatly impacted by distance, time or even some UV treatment, but are curbed by the amount of moisture in the air.

## INTRODUCTION

The COVID-19 pandemic has refocused attention on the need to understand and mitigate airborne transmission of viruses. Initially, transmission of SARS-CoV-2, the virus that causes COVID-19, was thought to occur mainly through coughing, person-to-person contact, and fomites (1). But, as our knowledge and understanding of both the virus and the disease increased, aerosol transmission emerged as the dominant form of SARS-CoV-2 spread (2–4). Airborne virus is believed to predominantly exist inside aerosolized particles of moisture emitted from the upper respiratory tract of infected persons through not only coughing, but speaking (3), singing (5), and even breathing (6). Coughing is known to produce larger, heavier droplets (diameter >10 μm) (7), while speaking, singing, and breathing are more likely to produce aerosols (defined here as diameter <10 μm) (3). Previous studies have shown that the duration of particle suspension in the air is a function of particle size; larger droplets fall quickly, whereas aerosols can remain suspended for long periods (8).

Indoor environments, such as gyms, restaurants, schools, or public transportation, have proven to be conducive to viral airborne transmission (5, 9–11). Data suggests that controlled ventilation (such as HVAC) can substantially lower transmission risk (9). Likewise, observations support the hypothesis that the risk of SARS-CoV-2 transmission is lower in school environments than in the general community when mitigation procedures, such as mask wearing, are strictly followed (12). In England, cases have been observed to increase with the return of students to in-person instruction (13). Even under stringent mitigation measures, in-school transmission was still found to be the cause of at least 5% of new cases among those in school buildings (12).

More recently, in-school mitigation protocols have been relaxed in many areas as case numbers have declined (14). This relaxation of mitigation measures may be problematic in the event of renewed infections since the majority of classrooms in US schools have insufficient levels of ventilation to provide for clean, safe indoor air (11, 15). Understanding the risk of transmission of not just SARS-CoV-2 but other airborne viruses in classrooms with no ventilation, therefore, is likely to be of high importance. Here, we report on an investigation into the ecology of unventilated classrooms using an aerosolized, airborne virus. We also introduce some mitigation strategies that may improve the health of the classroom’s occupants.

## METHODS

In a manner identical to Skanata and colleagues (16), we used a simple experimental setup to investigate potential asymptomatic aerosol transmission in built environments using a genetically modified *LacZ-****β***-marked phi6 bacteriophage as a proxy for SARS-CoV-2 (16–18). Phi6 is a lipid membraned phage that has similar size, shape, and physiological characteristics to SARS-CoV-2, but is not pathogenic and cannot infect humans, allowing for testing in built environments.

### Preparation of Viral Surrogate and Host

A single colony of LacZ-**α** producing *Pseudomonas phaseolicola* (18) was added to 30mL of lysogeny broth (LB) containing 30 mg of ampicillin and incubated for 18 h with rotary shaking (220 rpm) at 25 °C to produce an overnight culture for soft agar overlay plates. A bacterial lawn was created using 200 μL of overnight culture in 3 mL of soft agar to serve as detectors of viral infection for the experiment. When *LacZ-****β***-marked phi6 is plated on LacZ-**α** producing *P. phaseolicola* (18) and grown on LB agar supplemented with X-Gal, the resulting plaques appear blue. This use of *LacZ* complementation between phage and host ensured that any plaques observed on test plates resulted from aerosolized phage and not from naturally occurring phage.

To prepare our phi6 experimental lysate, 5 mL of stationary-phase *P. phaseolicola* overnight culture was added to 200 mL fresh LB. When the culture reached exponential phase,10 μL of frozen phage stock was added. Following 18 h incubation with shaking, phages were isolated by filtration through 0.22 μm syringe filters (Durapore; Millipore, Bedford, MA). Phage particles per mL (titer) were quantified via serial dilution and plating according to standard methods (19).

### Generation of Aerosols

The experiments used medical delivery nebulizers (UniHEART nebulizer, Westmed, Inc, Tucson, AZ) connected to an air compressor (Model 0399, Westmed, Inc, Tucson, AZ) to generate aerosolized viral droplets (MMAD 2-3 μm) at a rate of 6 L/min, similar to human respired volumes (20). To introduce phi6 into a classroom environment, 10 mL of the viral lysate was diluted in LB and inserted into the nebulizer. The diluted phage lysates had a concentration of approximately 10^8^ phage particles per mL (maximum viral load of experimental room per trial = 10 mL × 10^8^ phage/mL = 10^9^ phage per trial).

### Classroom Experiments in Non-ventilated Rooms

Standard petri plates containing LB agar supplemented with X-Gal and ampicillin were inoculated with the genetically modified strain of *P. phaseolicola* producing LacZ-***α***. These plates were placed vertically at varying distances from the nebulizer for up to 60 minutes of exposure. This design simulates a situation where an unmasked individual is spreading airborne viruses in a classroom. The plates were located 1, 1.8, 3.5, 5.5, and 7.3 meters (3, 6, 12, 18, and 24 feet respectively) away from the nebulizer to detect airborne transmission of phi6 (Figure S3 A & B). Two sets of four plates were placed on both the left and right sides with respect to the nebulizer at each of the different distances. Each plate within a set was exposed to the aerosolized phage for progressively longer durations of time for up to 60 minutes (i.e., 15, 30, 45, and 60 min). Traffic within the rooms was limited to the experimenter and only at the end of each 15 minute interval in order to cover petri plates as needed. Experimental trials were repeated over multiple days in two different non-ventilated rooms of the same dimensions at CUNY Queens College. The test rooms were located on different floors of the same building. Experiments were conducted on days with low humidity (<40% relative humidity) and high humidity (>40% relative humidity) under naturally occurring temperature conditions. Room temperature and RH were continuously monitored using wireless sensors (Model H5074, Govee, Shenzhen, China).

### Classroom Experiments with UV

An identical experimental setup was conducted in the same non-ventilated rooms with the presence of portable UV fan units (Arc Air, R-Zero, Model: RXAIR) supplied by CUNY Queens College. These UV units were designed to take in and disinfect air in occupied rooms, as opposed to units that use more powerful UV light sources to disinfect air and surfaces in unoccupied rooms. Air is brought into the unit by fans, in a manner similar to HEPA or Corsi-Rosenthal Boxes (21). For UV trials, one of the two nearly identical test rooms had two UV units, placed diagonally across from each other in the two corners of the room. The other room served as the control condition. In subsequent trials, the experimental room was used as a control room and *vice versa*. Data was collected under both low humidity (<40% relative humidity) and high humidity (>40% relative humidity) conditions. The UV units have 16 fan speeds. We tested at least three replicates each in the low (fan speed = 4), medium (fan speed = 8), and high (fan speed = 12) fan speeds.

### Experimental Titer

Plates were collected at the end of the experiment and placed in an incubator at 25°C. The volume of lysate remaining in the nebulizer at the end of the experimental trial was collected to determine the total volume of phage aerosolized. Titer of the experimental lysate was confirmed via plating. In 24 to 48 hours, infection of the *Pseudomonas* bacteria by marked phi6 in aerosolized particles resulted in blue plaques on the X-Gal containing medium (22).

## RESULTS

Results were found to be consistent with a previous study employing the same basic design (16), even without the mechanical ventilation used in that study. The relative exposure rate reported here was normalized to 4 × 10^7^ total phage released to account for any variation in the amount of phage aerosolized per trial. This normalized viral load is within the range of reported observations for SARS-CoV-2 (23). We define relative exposure rate as the rate of exposure per unit time (in 15-minute blocks) normalized to 4 × 10^7^ total phage aerosolized at any given distance. Exposure is quantified by counting the number of viral plaques which represent successful aerosolization, airborne transmission, and subsequent infection by phi6 into a bacterial host.

Our results indicate that there was a statistically significant increase in the exposure rates when relative humidity (RH) was below 40% when compared to RH higher than 40% (p value = 5.45 × 10^−7^, two-tailed T-test. Although plates closer to the sources at 1 and 1.8 m (3 and 6 ft, respectively) had higher overall exposure rates compared to the plates further away from the virus source, no additional effect by distance was observed. Instead, a strong and consistent trend of exposure rates dropping at RH greater than 40% was observed, even up to 7.3 m (24 ft) away from the source.

The same two laboratory classrooms were used for investigating the effect of commercially available portable UV air disinfection units, with one room acting as a control and one containing the UV fan unit(s) (Arc Air, R-Zero). In rooms with UV treatment, we did not see a statistically significant impact of UV disinfection on the mean number of plaques observed during similar periods of exposure, regardless of the speed at which fans were operated (Fig. 2). In fact, there was a slightly higher mean number of plaques in the UV treated rooms compared to the control (Fig. 3), although this effect was not statistically significant.

**Figure 1:**
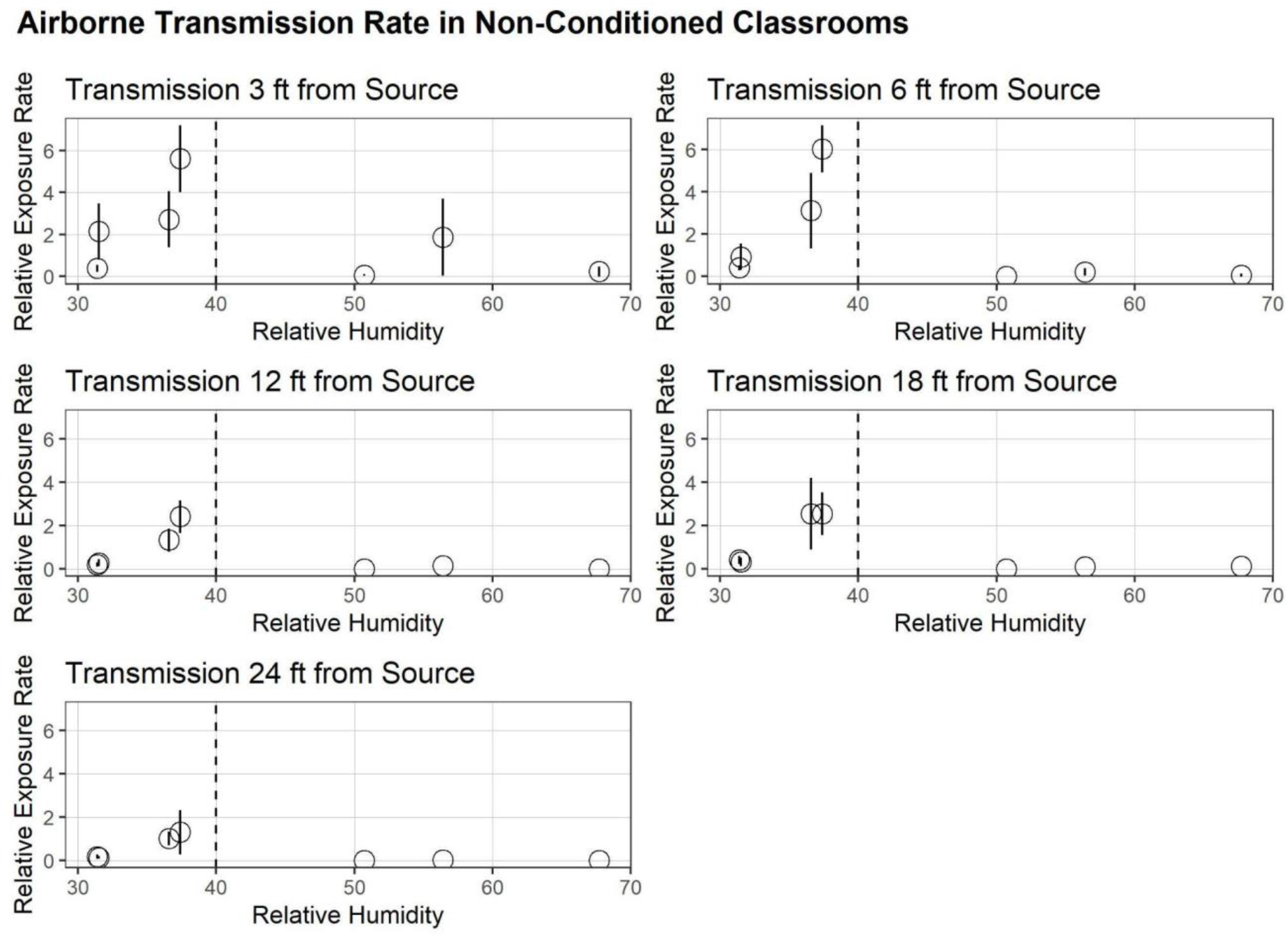
Relative exposure rates represent the rate of exposure per unit time (in 15-minute blocks) normalized to 4× 10^7^ total phage aerosolized in order to allow for comparison of multiple trials with variability in total number of phages aerosolized. At all distances we observe a drop off in the relative exposure rate when RH > 40%. Relative exposure rates are empirically higher within 2 m (6 ft) of viral source. Error bars represent 95% confidence intervals.

**Figure 2:**
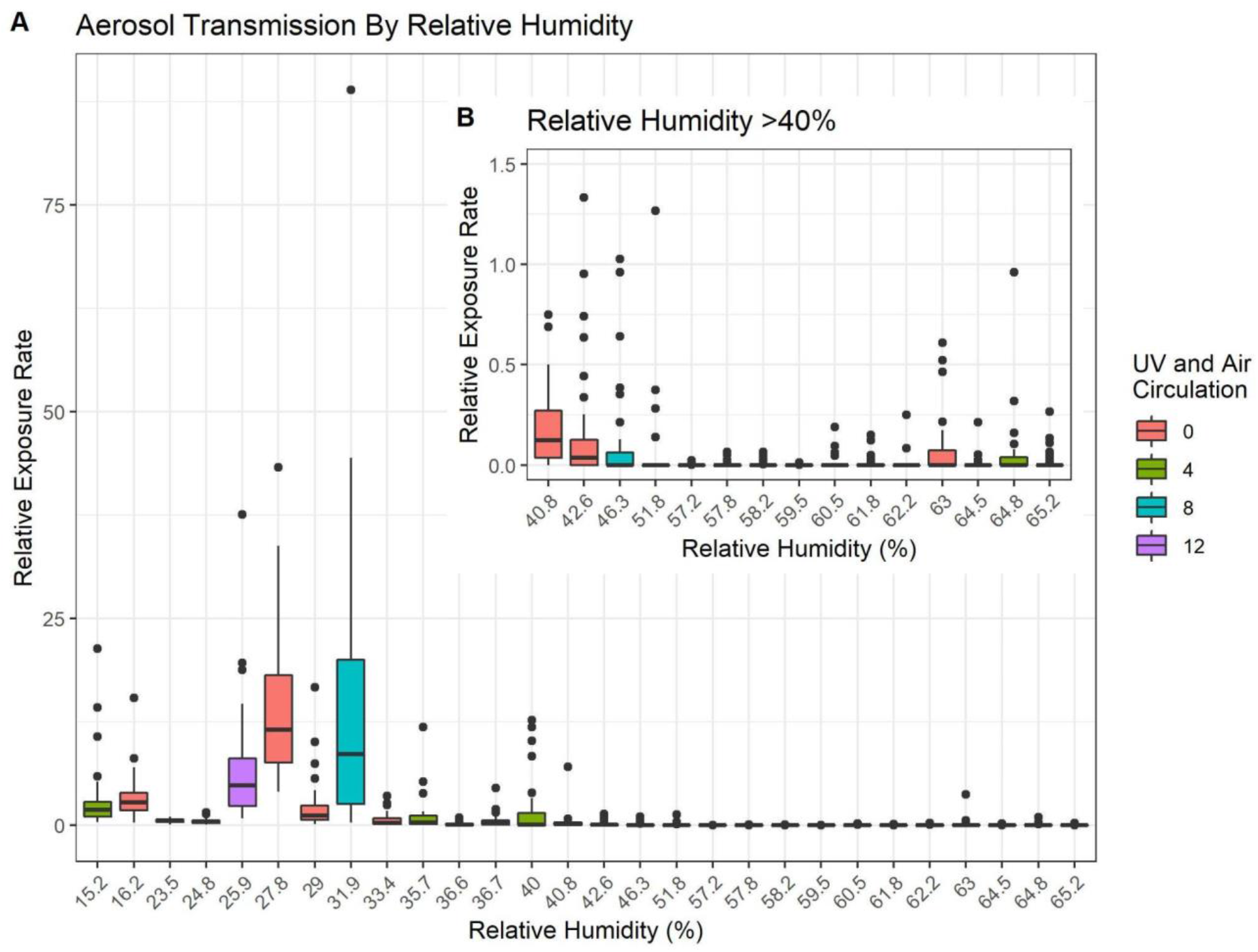
Relative Exposure Rates for all Trials. A. The relative exposure rates for all trials are arranged by RH. A drop in transmission is evident from RH ≥ 33.4%, however, there is a high degree of variability until RH ≥ 40.8%. B. The inset panel represents all trials with RH > 40% with the y-scale changed for easier viewing. Color indicates any UV treatment/fan speed. Red bars are no UV control conditions, green bars are UV treatment with “low” fan settings, blue bars represent UV treatment with “medium” fan setting, and purple bars show UV trials with fans set to “high” speed.

**Figure 3:**
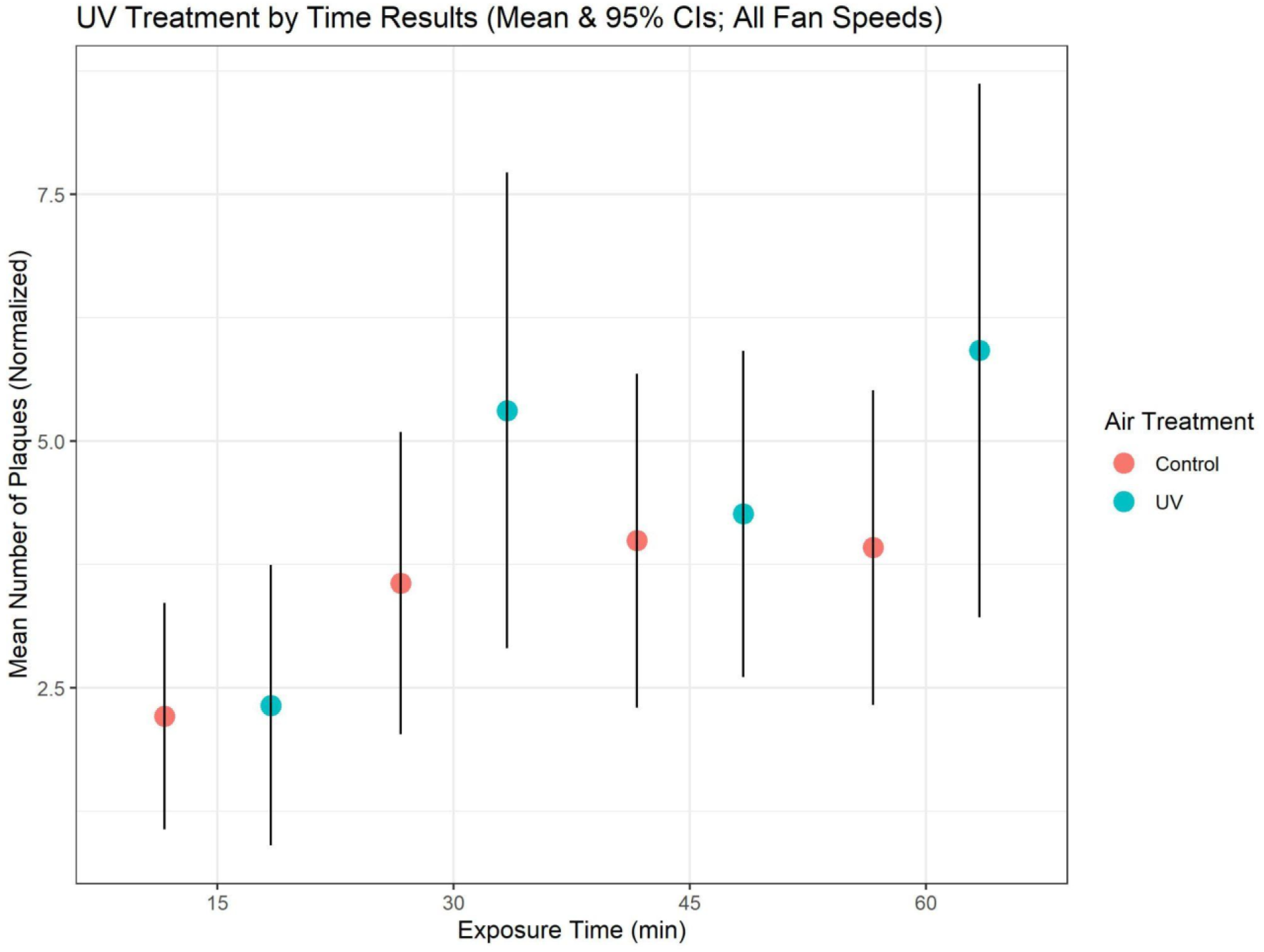
Mean number of phage plaques (normalized) by time of exposure and UV treatment. Attempts to disinfect the classroom air through use of portable UV-C generating devices had no effect upon the number of transmission events observed, regardless of the UV device fan speed or the amount of time of UV treatment. Some evidence suggests that the specific type of airborne virus as well as the wavelength of UV light along with various environmental conditions could affect disinfection results (26), although no impact was observed with the units used in this study.

Across all fan speeds, relative humidity remains the most significant factor observed. RH over 40% within UV trials have relative exposure rates below 1.5 on average. The maximum exposure rate within this group is 4, which is still extremely low compared to the exposure rates of RHs below 40%.

## DISCUSSION

Our results support the hypothesis that aerosolized viruses can and do transmit in indoor environments over long distances (up to 7.5 m or 24 ft) within minutes of their release, although the overall number of transmission events was reduced by half over such distances. This result is important in planning for any mitigation of airborne infectious diseases in the built environment in which humans spend a majority of our time (24). In particular, classrooms pose a potential risk as the people present spend long periods of time in these rooms, often without mechanical ventilation systems or sources of fresh air (25). Mitigation strategies to limit indoor transmission, then, become important, particularly in environments where there are low rates of air exchange.

We find that the most consistent environmental mitigation involves maintaining relative humidity above 40%. This RH level is not extreme or even uncomfortable for human or animal inhabitants. In addition, equipment capable of achieving and maintaining a RH of over 40% is not particularly expensive or difficult to install and run and can be easily added to a room environment without greatly impacting the designated activity in the room. For instance, in classrooms, multiple small or large humidifiers can be added to raise RH. In seasonal climates, warmer months are typically associated with higher RHs naturally. In cooler seasons, artificial (and modular) humidifiers can be put in place for a fraction of the investment cost required to upgrade mechanical ventilation.

Additional potential *ad hoc* mitigation measures include using UV-C light as a disinfectant. In our testing *in situ*, a portable commercially available UV disinfection unit did not perform as well as RH, even when multiple units were used in the same room. This result remained consistent regardless of fan speeds tested. No fan speed proved to be more effective than the control room tested contemporaneously without the UV units. Importantly, results observed for UV mitigation here may not be representative of all UV-based disinfection approaches or their efficacy, as device design and application vary widely (26).

## CONCLUSIONS

Mechanical ventilation standards for schools require minimum rates of fresh air per occupant per second (27). However, most schools in the United States fall far short of the minimum standards for mechanical ventilation, with associated losses in student performance (25). As such, low-cost mitigation techniques are needed. Baseline understanding of transmission rates in school rooms without mechanical ventilation is necessary in designing effective mitigation strategies. Here, we investigated airborne viral transmission in non-ventilated classrooms.

Our investigation found that successful infection events were not dependent upon distance or total time of exposure, although longer exposure time did lead to higher variability in transmission (Fig needed?). Instead, the strongest factor we observed in the rate of transmission was relative humidity.

Our data indicate that controlling RH in classroom environments is an effective and achievable mechanism to mitigating airborne viral transmission. In addition, humidity management and monitoring are low cost and can be done on an *ad hoc* basis in classroom environments, with little upfront costs or loss of space utilization. The threshold indicated for effective airborne pathogen mitigation is 40%, which is within the range of comfortable indoor environments for people, plants, animals, and equipment. Portable humidifiers can be used to help achieve the 40% RH target, with the added benefit that they can be shut down or moved when not in use. As a mitigation strategy, RH looks to be broadly applicable, low-cost, easily applied, and highly effective for classroom environments.

## ACKNOWLEDGEMENTS

We are grateful to Zeco Krcic and CUNY Queens College Buildings and Grounds in arranging for classrooms to be available for experimental trials and for supplying the UV units tested. We thank Sherin Kannoly for technical expertise. This work was funded by NSF RAPID Award No. 2032634 (JJD).

