## Supplemental Information for "Risk of Infection Due to Airborne Virus in Classroom Environments Lacking Mechanical Ventilation"

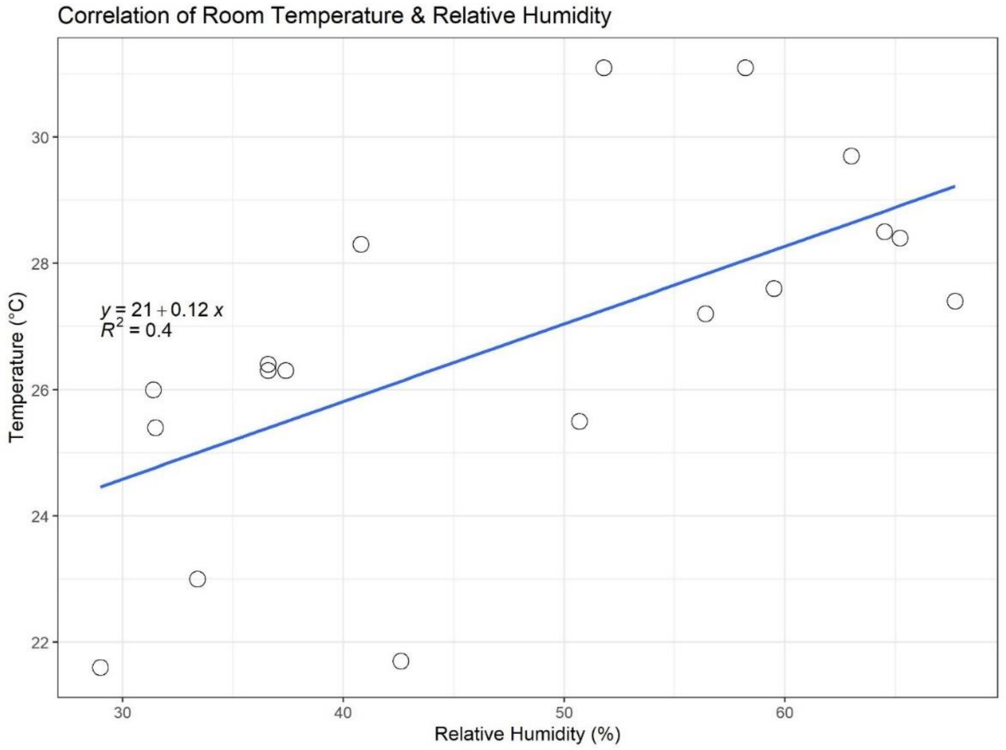


Figure S1: Correlation between Room Temperature and Relative Humidity: The average room temperature and RH are reported for all experimental and control trials. While RH generally correlated with temperature, the correlation was not strong ($R^2$= 0.4). In general, RH increased as temperature increased. This pattern is typical in most climates. The effect on transmission was found to be a result of RH and not temperature (see Fig. S2).

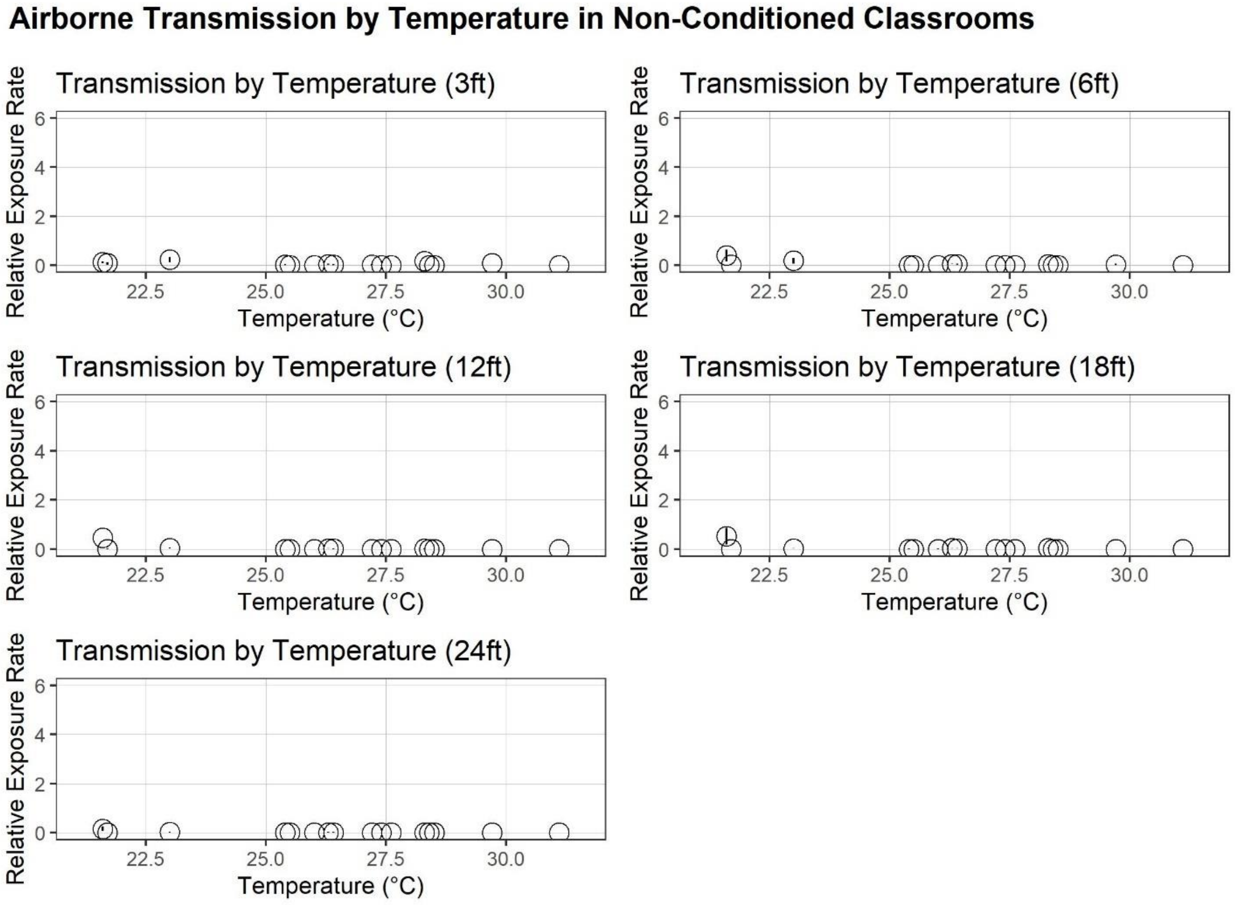


Figure S2: Transmission by Temperature at Each Distance Tested: Average room temperature did not show a correlation with relative exposure rate. Note that scale of Y-axis is set to match that in Figure 1.

A

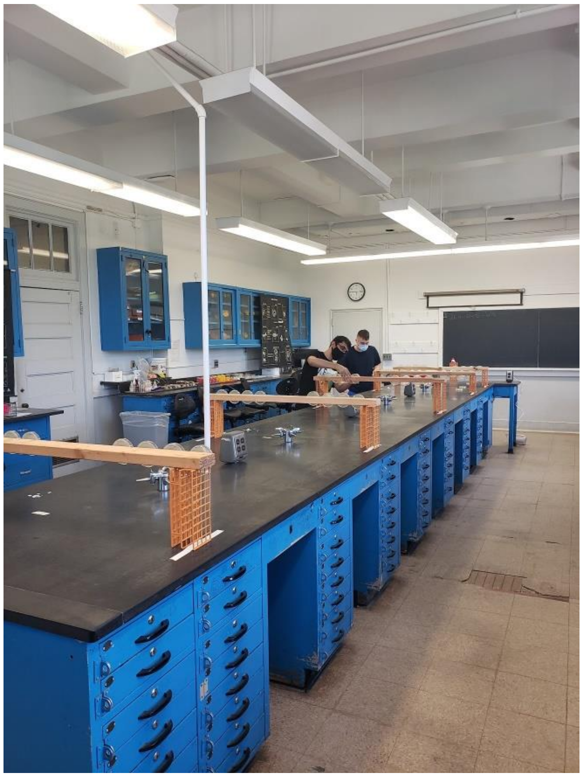

B

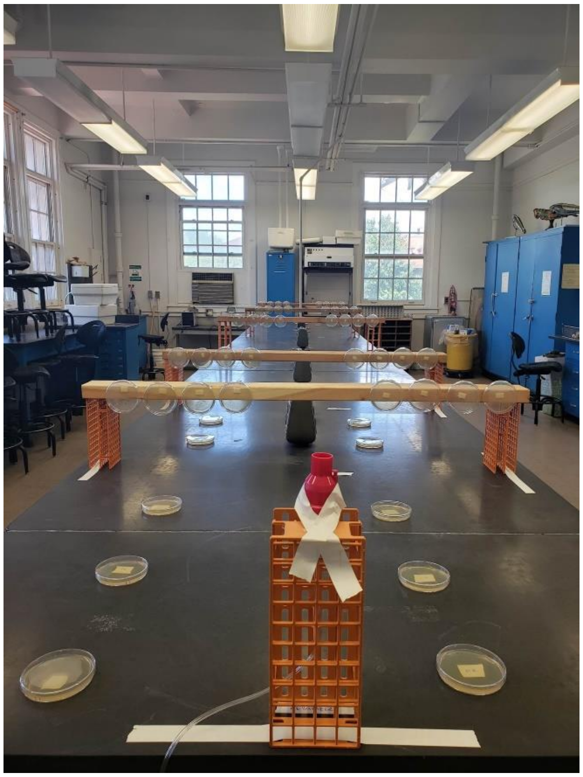

Figure S3: Experimental Setup in Classrooms Without Mechanical Ventilation. A. Experimenters set up petri plates seeded with lawns of LacZ-**α** producing *P. phaseolicola* on LB agar. Plates were installed vertically on both the left and right sides of the room in rows of 4 on each side, one plate for each block of timed exposure (15, 30, 45, and 60 minutes). B. A medical-grade nebulizer was the source of generated aerosols laden with *LacZ-**β***-marked phi6 bacteriophage (pink device, center foreground). Vertical detector plates were arranged on left and right sides of the nebulizer at 1, 1.8, 3.5, 5.5, and 7.3 meters (3, 6, 12, 18, and 24 feet respectively). Horizontal plates were used near the nebulizer to verify large droplets falling out of the air.
